# Mass lysis of bacterial predators drives the enrichment of antibiotic resistance in soil microbial communities

**DOI:** 10.1101/2023.11.20.567171

**Authors:** Saheli Saha, Jyotsna Kalathera, Thoniparambil Sunil Sumi, Vishwadeep Mane, Sina Zimmermann, Silvio Waschina, Samay Pande

**Affiliations:** Bacterial Ecology and Evolution Group, Department of Microbiology and Cell Biology, Indian Institute of Science, Bangalore, India; Institute of Human Nutrition and Food Science, Nutriinformatics, Kiel University, Kiel, Germany

## Abstract

While studies on anthropogenic activities and antibiotic resistance are numerous, the impact of microbial interactions on resistance in complex communities remains uncertain. Here we demonstrate a correlation between the presence of *Myxococcus xanthus* in natural soil communities and the abundance of antibiotic-resistant bacteria. Further, introducing *M. xanthus* isolates also enriches antibiotic resistance. This is due to the mass lysis of *M. xanthus* cells, which results in a toxic environment that fosters the proliferation of pre-existing resistant bacteria rather than de novo resistance evolution. Metagenomic analysis revealed that this enrichment is not limited to the tested antibiotics in culture-based methods, indicating its broader relevance. Crucially, these findings go beyond laboratory settings, showing *M. xanthus* introduction enriches resistant isolates in natural soil communities. Finally, we demonstrate that the mass lysis of *M. xanthus* cells during starvation-induced development—key aspect of the lifecycle of *M. xanthus*—also results in the enrichment of antibiotic resistance in soil communities. Together, we demonstrate how life-history traits in bacterial predators, like *M. xanthus*, significantly impact antibiotic resistomes in nature. This study also highlights the complex dynamics at play in the evolution and maintenance of antibiotic resistance, emphasizing the role of interspecies interactions in shaping antibiotic resistance profiles.

## Introduction

Advances in sequencing and molecular technologies over last decade have revealed a previously unfathomable distribution and abundance of antibiotic resistance in varying geographical^1–4^ and ecological^5–7^ conditions. Anthropogenic activities such as rampant use of antibiotics and husbandry have been attributed to the evolution of novel resistant mechanisms as well as their distribution^8–10^. In addition to rapid emergence of resistance in environments with extreme antibiotic concentrations, resistance can also emerge and be maintained for longer duration in extremely low concentrations of antibiotics^11–14^. Thus, natural antibiotic resistome is likely affected by the antibiotics irrespective of whether their concentration and source.

Bacteria use diverse mechanisms to antagonise each other, including production of antibiotics. A number of microbial species are proficient antibiotic producers^15–18^. Such microbes generally harbor multiple biosynthetic clusters and show ability to produce distinct antibiotics^19–23^. However, influence of antibiotic producing microbes on the emergence or maintenance of resistance mechanisms in nature as well as in lab environment remains little explored.

*Myxococcus xanthus* is a ubiquitous^24–26^ soil living gram negative predatory bacteria that forms multicellular spore bearing fruiting bodies upon starvation^27^. During vegetative growth *M. xanthus* can feed on freely available nutrients as well as on other microbes. Killing of prey bacteria by *M. xanthus* is mediated by both contact dependent and independent mechanisms such as antibiotic production^17,28^, secretion of toxins^29^, lytic enzymes^30^, and secretion systems^31^. We hypothesised that the expression of antagonistic traits especially antibiotic production might influence both physiological as well as evolutionary response by local microbial community.

Here, we show that the presence or introduction of *M. xanthus* in soil communities both in natural as well as lab environment can result in enrichment of antibiotic resistance. Interestingly, changes in the frequency of resistant isolates is brought about by rapid death of *M. xanthus* population in our microcosm experiments. Moreover, our results demonstrate that starvation induced death of *M. xanthus* results in the release of substances that are growth inhibitory to non-myxobacterial species and hence can result in the enrichment of bacteria resistant to the growth inhibitory substances released by dying *M. xanthus* cells. Together, we demonstrate a unique instance of single species of bacteria affecting overall maintenance of antibiotic resistance in complex microbial communities.

## Results

### Presence of *M. xanthus* results in enrichment of antibiotic resistant isolates in natural soil communities

We tested whether presence of *M. xanthus* is correlated with either increased or decreased abundance of antibiotic resistance in natural soil communities. To do so, twenty-five random soil samples were first categorised as the ones in which *M. xanthus* was detected (16 out of 25) and the ones in which *M. xanthus* was not detected (9 out of 25). These samples were named as *M. xanthus* positive and *M. xanthus* negative soil samples, respectively. Next, frequencies of non-myxobacterial isolates resistant against clinically relevant antibiotics (Ampicillin, Gentamycin, Kanamycin, Rifampicin, Tetracycline, Vancomycin) were measured. These experiments revealed that the mean frequency of antibiotic resistant isolates in *M. xanthus* positive samples (0.5878 %) was higher relative to the samples in which *M. xanthus* was not detected (0.0012 %) (Figure 1A, independent-sample t-test for differences between arcsine square root transformed frequency of resistant microbes in *M. xanthus* positive soil and *M. xanthus* negative soil sample p = *1.941e^−14^*).

**Figure 1.**
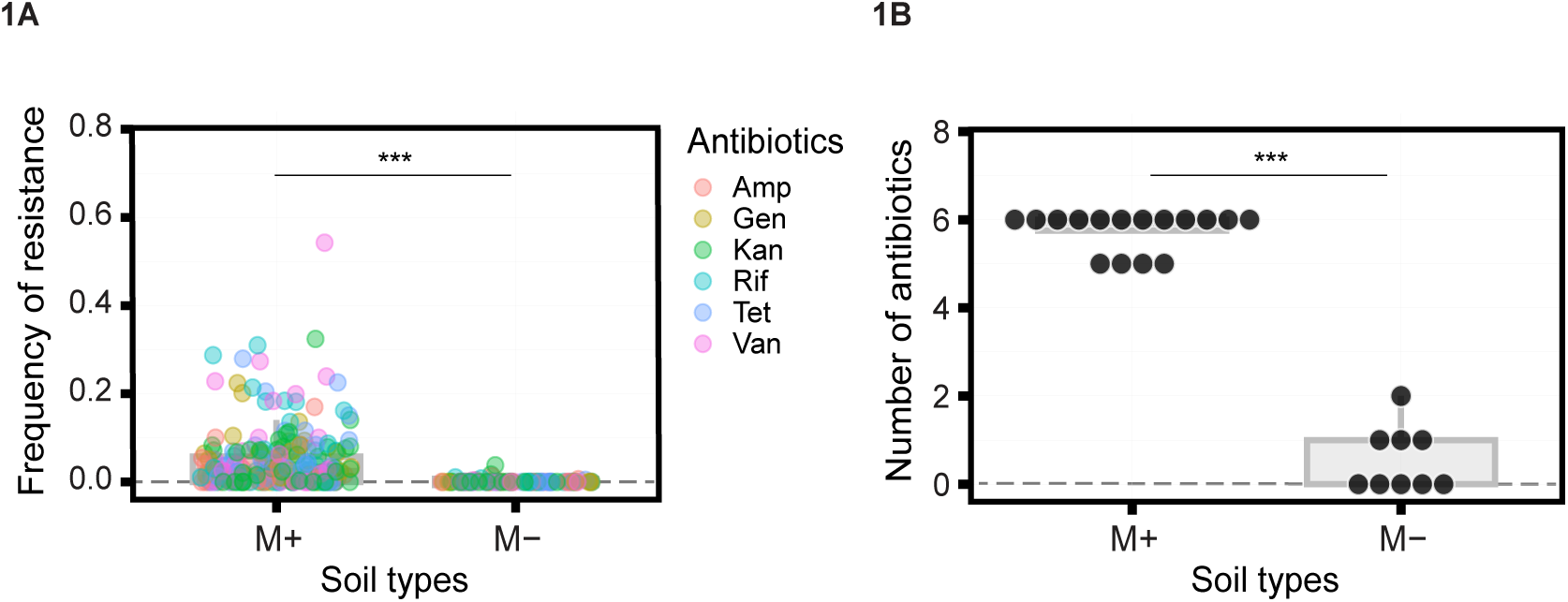
Higher frequency of antibiotic resistant bacteria is correlated with presence of *M. xanthus* in natural soil samples. A) Overall frequency of antibiotic resistant isolates across twenty-five distinct locations and three distinct media types is shown. Independent-sample t-test for the differences in the arcsine square root transformed frequency of antibiotics resistant isolates between soil sample in which *M. xanthus* was detected (M+) (16 out of 25) and the ones in which *M. xanthus* was not detected (M-) (9 out of 25), p = *1.941e^−14^*. Each dot represents distinct soil community and different colours represent different antibiotics (Ampicillin, Gentamycin, Kanamycin, Rifampicin, Tetracycline, Vancomycin). B) Number of antibiotics against which resistant isolates were found in individual soil sample is shown. Independent-sample t-test for differences between soil samples in which *M. xanthus* was detected (M+) (16 out of 25) and the ones in which *M. xanthus* was not detected (9 out of 25) (M-), p = *0.0000656*. Each dot represents distinct soil community and different colours represent different antibiotics (Ampicillin, Gentamycin, Kanamycin, Rifampicin, Tetracycline, Vancomycin).

Higher frequencies of resistant isolates can be the result of either one or many distinct isolates carrying resistance against a minority of the antibiotics used in our study. However, we observed resistance against majority of antibiotics in *M. xanthus* associated communities (Figure 1B, independent-sample t-test for differences between soil samples in which *M. xanthus* was detected and the ones in which *M. xanthus* was not detected, p = *0.0000656*). Twelve out of sixteen *M. xanthus* positive soil samples harbored resistance against each of the six antibiotics and remaining four samples harbored resistance against five out of six antibiotics used. Whereas similar analysis of *M. xanthus* negative soil did not exhibit such high prevalence of resistance against the diversity of antibiotics tested.

Abundance of antibiotic resistant isolates was estimated using three distinct media types (see methods). Therefore, we also tested whether the media types in addition to the presence or absence of *M. xanthus* can influence the frequency of antibiotic resistant isolates (Figure S2, independent-sample t-test between M+ and M-conditions in each media type, LB (0.1X): p = *3.864e^−12^*; LB (1X) : p = *2.2e^−16^*; TSA (0.1X) : p = *1.002e^−08^*). Our analysis revealed that only presence or absence of *M. xanthus* had an effect on the resistance frequency and neither the media nor the identity of the soil community had an effect. Further, relative frequency of resistant isolates for different antibiotics varied greatly among distinct soil samples (Figure S1). Together, these results show correlation between presence of *M. xanthus* and enrichment of antibiotic resistance in nature.

Strong correlation between the presence of *M. xanthus* with a higher antibiotic resistance frequency in natural soil communities suggested that *M. xanthus* is responsible for the abundance of antibiotic resistant bacteria. To test this hypothesis, the effect of addition of *M. xanthus* on the changes in the abundance of antibiotic resistant isolates was measured. For this, *M. xanthus* was inoculated with four randomly sampled soil communities in an antibiotic free environment, and frequency of antibiotic resistant bacteria was estimated after 6 days of co-culture. These experiments revealed that across all the four soil communities introduction of *M. xanthus* resulted in an enrichment of resistant isolates across six clinically relevant antibiotics and three media conditions. On an average, across four soil communities 0.902 % isolates exhibited resistance to at least one or more antibiotics tested when the consortia were co-inoculated with *M. xanthus*. Whereas mean frequency of antibiotic resistant isolates was significantly lower (0.000027 %) in soil communities which were not co-inoculated with *M. xanthus* (Figure 2A, paired-sample t-test for differences between percentage of resistant isolates when soil communities were cocultured with or without *M. xanthus*, p < *0.000004* and Figure S3A, paired-sample t-test between M+ and M-conditions in each media type, LB (0.1X): p = *4.218e^−07^*; LB (1X) : p = *5.554e^−07^*; TSA (0.1X) : p = *4.467e^−05^* )

**Figure 2.**
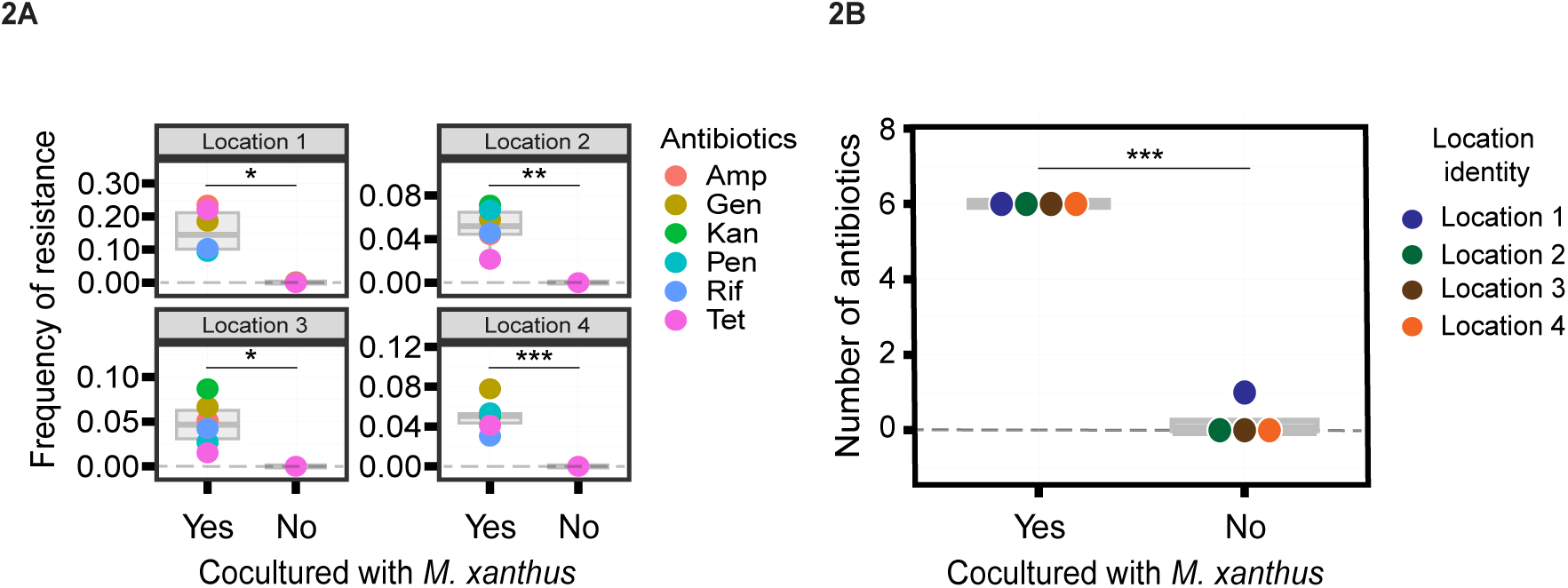
Antibiotic resistant isolates are enriched in soil communities cocultured with *M. xanthus*. Soil from four distinct locations was cultured either with or without *M. xanthus*. A) Frequency of resistant isolates post 6 days of incubation are shown. Paired-sample t-test for the differences in the arcsine square root transformed frequency of antibiotics resistant isolates between soil sample in which *M. xanthus* was added and the ones in which *M. xanthus* was not added, p < *0.000004*. Each dot represents distinct soil community and different colours represent different antibiotics (Ampicillin, Gentamycin, Kanamycin, Rifampicin, Tetracycline, Vancomycin) B) Number of antibiotics against which resistant isolates were found is shown. Paired-sample t-test for differences in the number of antibiotics against which resistance was observed in the two experimental conditions, where *M. xanthus* was added or not added, p = *0.00018*. Each dot represents distinct soil community and different colours indicate different locations.

We tested whether the enrichment of antibiotic resistant microbes brought about by *M. xanthus* is a more generalised response to presence of any *M. xanthus* isolate, or it is specific to the isolate used here. To do so, we repeated the assay with six randomly selected natural isolates of *M. xanthus*. These experiments revealed that only three of the six natural isolates tested could influence frequency of antibiotic resistant microbes in soil communities (Figure S3B, paired-sample t-test between day 0 and day 6 post addition of each *M. xanthus* to the community, p < *0.05*). These results demonstrate that not all *M. xanthus* isolates can have the similar effect on the frequency of antibiotic resistant bacteria as the focal strain S3 used in this study.

Although introduction of *M. xanthus* resulted in enrichment of antibiotic resistant isolates, to further understand the dynamics of increase in the frequency of antibiotic resistant bacteria upon introduction of *M. xanthus*, we simultaneously tracked both *M. xanthus’s* growth and overall resistance frequency over the course of co-culture experiment. For this, four distinct soil communities were cocultured with *M. xanthus*, and population size of *M. xanthus* and frequency of antibiotic resistant bacteria was measured every day for 6 days. These experiments demonstrated that *M. xanthus* could grow and increase in number for the first day, and then showed a drastic population decline reaching below detection limit after 2 days on incubation (Figure 3A ). Further, the frequency of resistance isolates increased only after day 2 of incubation i.e. once *M. xanthus* could no longer be detected. These findings suggest that the death of *M. xanthus* might be responsible for the enrichment of antimicrobial resistance in soil communities.

**Figure 3.**
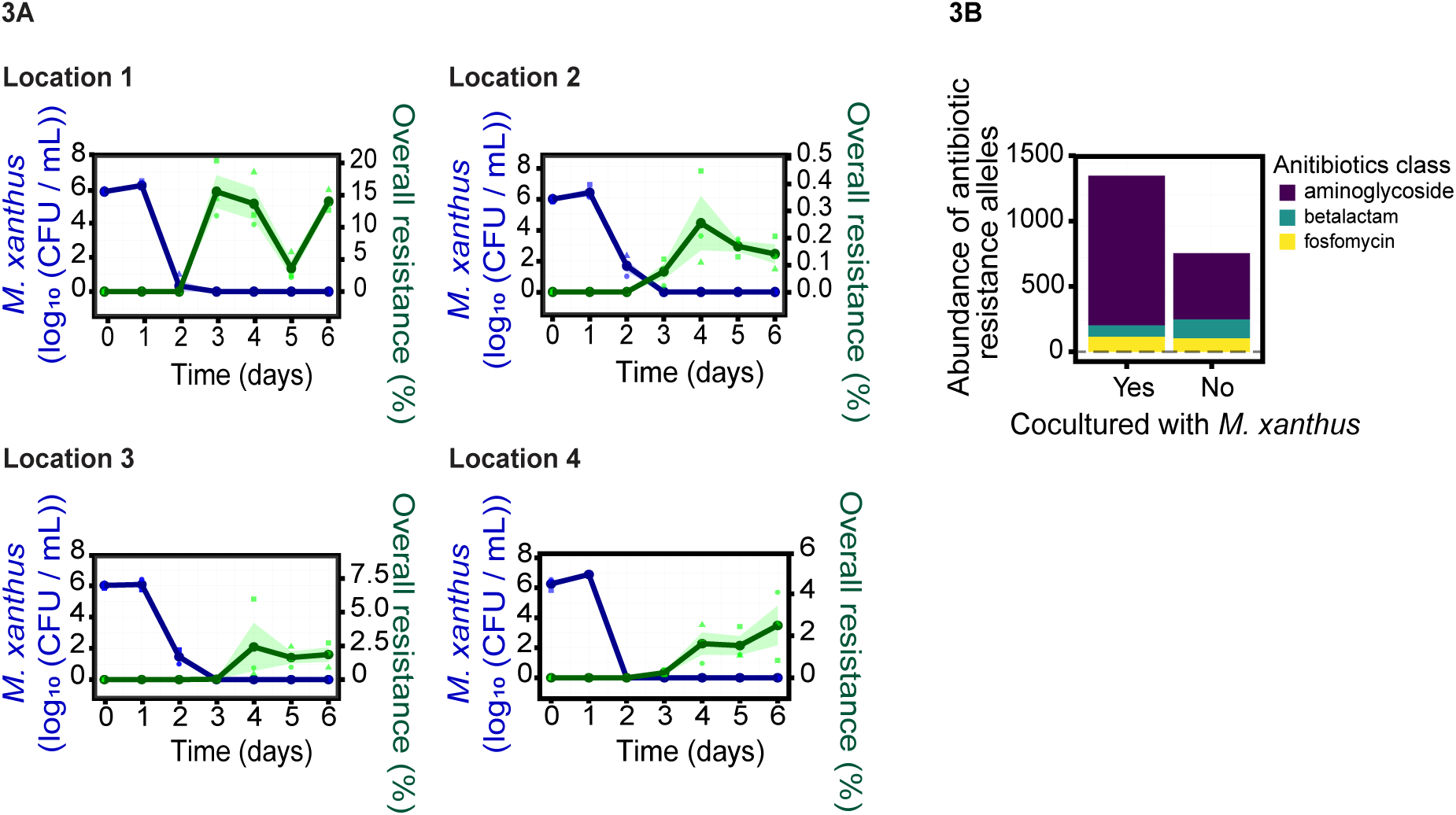
Frequency of antibiotic resistant bacteria increases only after the death of *M. xanthus* population. A) Green lines represent changes in overall percentage of resistant isolates over 6 days, in four distinct soil communities when cocultured with *M. xanthus*. The concurrent *M. xanthus* cell counts (log (CFU/ml)) in all four communities are represented by the blue lines. Squares, triangles, and circles in lighter colours represents average frequency of antibiotic resistant bacteria (for six antibiotics i.e., Ampicillin, Gentamycin, Kanamycin, Rifampicin, Tetracycline and Vancomycin) within respective independent replicate (n = 3). B) Abundance of the antibiotic resistance alleles were determined from metagenome data obtained from second location using KmerResistance algorithm. The abundance of various resistance class alleles when soil communities were either cocultured with *M. xanthus* or grown alone for 6 days is shown. Different colours show the distinct classes of antibiotic resistance alleles.

Our culture based detection method can detect frequency of resistant isolates against the antibiotics used in the assay, and may not reveal the overall abundance of antibiotic resistance across distinct antibiotic types. Therefore, to detect if addition of *M. xanthus* can alter the overall abundance of antibiotic resistance alleles within soil communities, total DNA was extracted from the communities that were either cocultured with *M. xanthus* or grown in its absence. Shotgun metagenome analysis of the extracted DNA revealed that antibiotic resistance genes were indeed enriched in the communities that were cocultured with *M. xanthus* (Figure 3B), compared to when *M. xanthus* was not added (Figure 3B). Moreover, amongst the three major classes of antibiotic resistance genes (aminoglycosides, beta lactams and fosfomycins) that were detected in these communities, resitance against aminoglycosides enriched the most in the communities that were cocultured with *M. xanthus* (Figure 3B). Together, metagenome analysis supports the results from culture based assays.

### Mass-lysis in *M. xanthus* population can modulate the environmental toxicity resulting in enrichment of antibiotic resistance

Since, bacteria are known to release antibiotics and growth inhibitors upon cell death^32–35^, we hypothesised that mass-lysis of *M. xanthus* population might also result in release of growth inhibitory diffusible substances in environment. *M. xanthus* is known to have a large repertoire of secondary metabolites that contributes differentially to specific life-cycle stages. For example, secretion of antibiotics, toxins, and enzymes during predation, secondary metabolites during development^36^ and diffusible substances to help germination^37^. Sudden release of growth inhibitory substances such as antibiotics and toxins because of mass lysis of *M. xanthus* might results in enrichment of microbes resistant to such substances.

To test this hypothesis, the spent media from coculture experiments in which *M. xanthus* was co-incubated with soil communities, were extracted at every 24 h interval. Freshly harvested soil communities were then inoculated in the medium conditioned with the harvested spent medium. If lysis of *M. xanthus* on day 2 (Figure 3A) results in the release of diffusible molecules that can potentially enrich antibiotic resistant bacteria, then addition of spent media from day 2 to soil communities should also result in the enrichment of antibiotic resistant bacteria. Whereas, spent media from day 1 and day 0 should not result in any enrichment. Since all of the results were consistent across six antibiotics used in earlier experiments, we measured resistance against three representative antibiotics from different classes. This was primarily done for the ease of logistics. In line with our expectations, addition of spent media from second day onwards can results in enrichment of resistance frequencies in complex soil communities (Figure 4A). We further show that the diffusible molecules present in the spent media from 2 day old coculture is indeed growth inhibitory to different lab strains of soil bacteria (*A. globiformis, E. coli, P. putida and R. vitis*) (Figure 4B, one-sample t-test [FDR corrected] for relative growth of respective species, p < *0.02*). Together, these result support the hypothesis that release of growth inhibitory substances in the environment by dying *M. xanthus*, may select for resistant isolates, thereby enriching overall antibiotic resistant isolates in complex communities.

**Figure 4.**
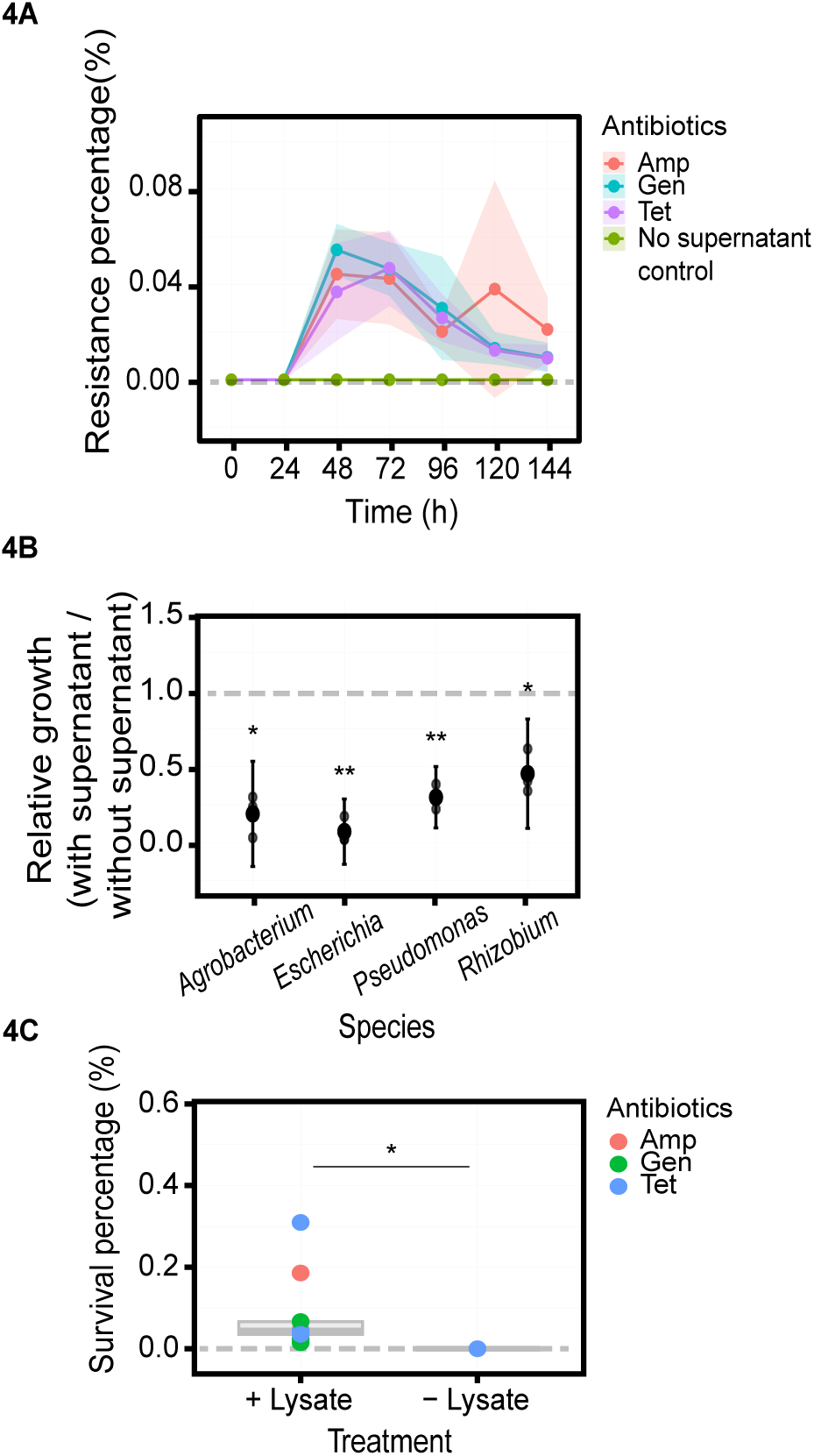
Lysis of *M. xanthus* cells is responsible for enrichment of resistant isolates in complex soil communities. A) Spent media from soil communities that were cocultured with *M. xanthus* results in enrichment of resistant isolates. Distinct lines represent the percentage of isolates resistant against three distinct antibiotics (Ampicillin, Gentamicin and Tetracycline). Green line at 0.00 is the control where no spent media was added to the soil communities. B) Spent media from soil communities that were cocultured with *M. xanthus* inhibits the growth of soil bacteria. Each dot represents growth of bacterial species in the presence of supernatant relative to their growth in the absence of supernatant. Error bars represents 95% confidence intervals. One-sample t-test [FDR corrected], to test if the relative growth is different from one, *p <*0 .05*, **p <*0.005* (n = 3). C) Introduction of supernatant extracted from *M. xanthus* lysate can enrich resistant isolates in complex soil communities. Percentage of resistant isolates when *M. xanthus* lysate is either added or not added is shown. Each dot represents distinct soil sample and different colours represent three distinct antibiotics (Ampicillin, Gentamicin and Tetracycline). Paired-sample t-test between percentage resistant isolates in the presence of lysate compared to in its absence, p = *0.032* (n = 3).

Although our results show that the enrichment in antibiotic resistance is most likely brought about by presence of diffusible growth inhibitory molecules in the environment, we further tested if these molecules are specifically released as a result of lysis of *M. xanthus* cells. To do so, *M. xanthus* cells were lysed by sonicating the vegetative cells (See methods), and the soil communities were cultured in the medium with and without *M. xanthus* lysate. These experiments demonstrated that on an average, introduction of *M. xanthus* lysate results in enrichment of antibiotic resistant isolates by 6472 percentage folds (Figure 4C, paired-sample t-test between resistance frequencies where lysate was added or not added, p = *0.032*). Identification of the nature of the active molecules released by lysis of *M. xanthus* requires extensive investigation. Hence, though it would be intriguing to identify the active molecules responsible for these observation, this aspect of the study is part of future investigations.

### Enrichment of AMR in natural soil communities are most likely brought about by cell lysis during development phase of *M. xanthus*

Starvation induced development of spore filled fruiting body is an important aspect of *M. xanthus* biology. Importantly, during the development process majority of (∼ 90 %) *M. xanthus* cells die^38^. Thus, death of majority of population is part of *M. xanthus* lifecycle. We hypothesised, that in nature, death during development might be responsible for the release of growth inhibitory substances that enrich antibiotic resistance in soil (Figure S5). To test this, supernatant was extracted from population of *M. xanthus* after allowing them to form fruiting bodies on starvation media. Freshly extracted soil communities were then cultured either in the presence or the absence of this supernatant. Estimation of the frequency of antibiotic resistant bacteria in these experiments revealed that the addition of supernatant extracted from starved population of *M. xanthus* results in the enrichment of antibiotic resistant isolates by 13 % compared to when supernatant was not added (Figure 5B, Paired-sample Wilcoxon test, p = *0.048*). Hence, since mass-lysis of *M. xanthus* population is an essential part of its starvation response phase of the life cycle, abundance of antibiotic resistant isolates in the presence of M. xanthus might be because of the presence diffusible substances released by dying cells of *M. xanthus* during fruiting body formation.

**Figure 5.**
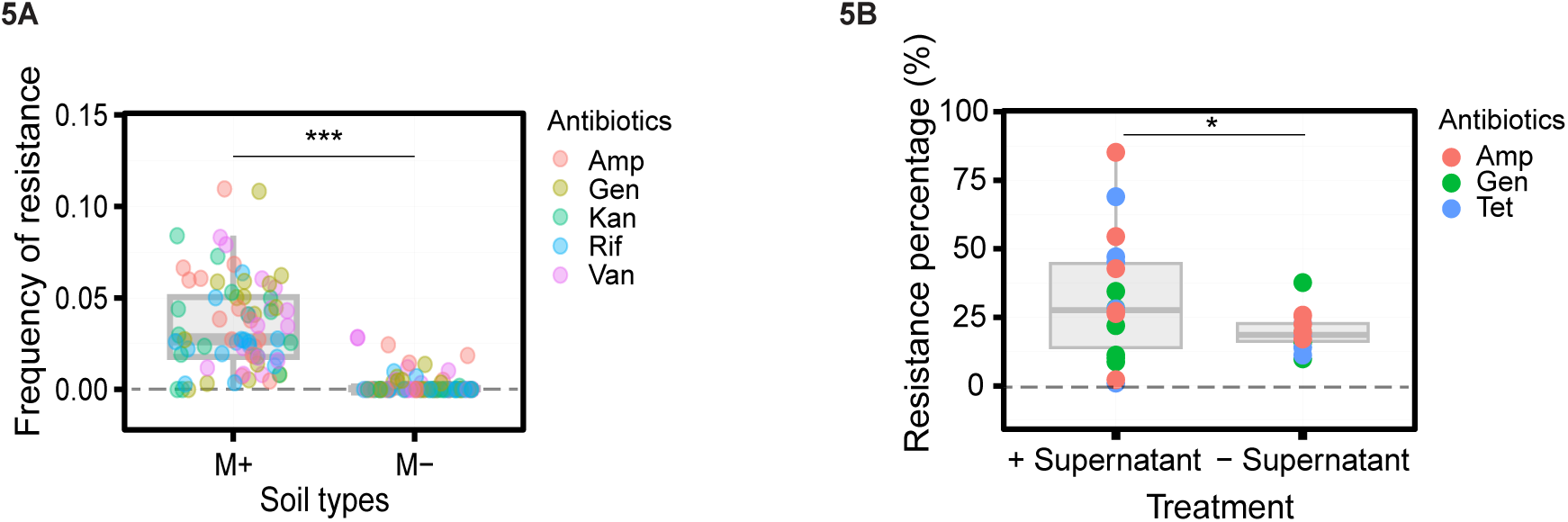
Mass-lysis of *M. xanthus* cells during fruiting body development results in enrichment of antibiotic resistant isolates in the environment. A) Soil communities cocultured with *M. xanthus* show enrichment of antibiotic resistance. Frequency of antibiotic resistant isolates in soil communities is shown. Paired-sample t-test for differences in the arcsine square root transformed frequency of resistant isolates in the presence of *M. xanthus* (M+) relative to in absence of *M. xanthus* (M-), p = < *2.2e^-16^* (n = 3). B) Addition of supernatant extracted from *M. xanthus* fruiting bodies to three distinct soil communities results in enrichment of antibiotic resistance. (Paired-sample Wilcoxon-test between the percentage resistances in conditions where supernatant was added or not added, p = *0.048* (n = 6).

Significance of reproducing laboratory experiments in nature has been highlighted time and again^39,40^. While our laboratory experiments were conducted on liquid minimal media, we understand that conditions in soil may vary significantly, both in terms of nutrient content as well as structuredness of the environment. Additionally, while laboratory controlled experiments allows the maintenance of a constant abiotic conditions, it is expected that in nature, fluctuations in abiotic conditions are more common. Since both spatial structure and abiotic fluctuations can significantly influence community structure and ecological interactions^41–44^ it was possible that our observations may not hold true in nature. Hence, we investigated whether the findings of increased antibiotic resistance upon introduction of *M. xanthus* in complex soil communities, in lab conditions can also be reproduced in natural soil environment. Interestingly, similar to the outcomes from the experiments performed in liquid medium in lab conditions, introduction of natural isolate of *M. xanthus* in soil pockets in natural locations also results in increased frequency of resistant isolates. Overall, samples with *M. xanthus* contained 0.186 % percent resistant isolates relative to 0.005 % percent in the samples where *M. xanthus* was not inoculated. Combined with the first set of results which showed correlation between abundance of resistance and presence of *M. xanthus*, these results demonstrate that in nature *M. xanthus* strongly modulates abundance of antibiotic resistant microbes (Figure 5A, paired-sample t-test for differences between arcsine square root transformed frequency of resistant microbes when soil communities were cocultured with *M. xanthus* and control p < *2.2e^-16^*).

## Discussion

Our results demonstrate that both in lab as well as natural conditions, resistance to clinically used antibiotics is enriched in the presence of *M. xanthus*. First, we observed that *M. xanthus* associated soil communities harbor significantly higher frequency of antibiotic resistant isolates relative to the communities that seem to be not associated with *M. xanthus*. Next, we demonstrated that introduction of *M. xanthus* in soil communities in lab as well as in natural conditions results in enrichment of antibiotic resistant isolates. Contrary to our expectations, antibiotic resistant isolates were not enriched directly by the presence of *M. xanthus* within the communities. Instead, the changes in the microbial communities detected in our study were manifested by death of *M. xanthus.* Taken together, we demonstrate that death of *M. xanthus* population, can result in the enrichment of antibiotic resistant bacteria. Further, the death of *M. xanthus* populations can be the result of the lifecycle of *M. xanthus* in which majority of the cells die during starvation induced fruiting body development, possibly explaining the reason for enrichment of the antibiotic resistant bacteria in the presence of *M. xanthus* in natural communities. Significantly, our result demonstrates the correlation between presence of *M. xanthus* and abundance of antibiotic resistant bacteria in the nature and explains the reason behind it.

The increased frequency of resistant isolates is most likely because of the enrichment of pre-existing resistant bacteria in the soil communities and not a result of de-novo evolution of resistance. This is because, all the soil samples used in our study exhibited antibiotic resistance even before the start of the co-inoculation experiments with *M. xanthus*. Enrichment in frequencies of antibiotic resistant bacteria in these communities can either result from the ability of the resistant isolates to grow in the toxic environment or by simply persisting in that environment, while the abundance of the sensitive isolates declines. This is also evident in the metagenome analysis which revealed presence of genetic determinants previously reported to be responsible for antibiotic resistance. Moreover, the experiments were conducted for a total duration of six days, which is a relatively shorter time span for de-novo evolution of resistance against variety of antibiotics to emerge and to raise to the frequencies observed in our experiments.

Our experiments were conducted using natural soil communities, which might explain large differences in the outcomes of the experiments. Further, we primarily report increased abundance of antibiotic resistant isolates among culturable microbes. Therefore, it is important to be cautious and refrain from extrapolating the results to complete microbial communities where unculturable microbes are a dominant fraction. However, the metagenome analysis does reveal that the observations are likely to be true for overall microbiome which includes unculturable majority. Importantly, since most bacterial pathogens are culturable microbes that grow on heterotrophic proteinaceous medium lend importance to the findings reported here, even if the effects shown here are limited to culturable heterotrophic microbial diversity.

Death of *M. xanthus* cells is responsible for the changes in the frequency of antibiotic resistant bacteria. Important aspect of *M. xanthus* biology, i.e., formation multicellular spore filled fruiting bodies upon starvation is associated with death of majority of the population of *M. xanthus*. Analysis of the sporulation efficiency of natural isolates of *M. xanthus* revealed that the highest sporulation efficiency among natural isolates was approximately ten percent^38^. Thus, at least ninety percent of vegetative cells in a population die during starvation induced development, and only minority become spores. Results presented in this manuscript demonstrate that the diffusible substances released during the fruiting body formation results in the enrichment of antibiotic resistant bacteria. Thus, we predict that in nature too, death of *M. xanthus* because of feast and famine cycles that drive growth and fruiting body development of *M. xanthus* respectively, will result in the enrichment of antibiotic resistant microbes.

Although, we show that it is indeed the lysate of *M. xanthus* that enriches resistance in prey, the precise mechanism or the active molecules that are primarily responsible for this environmental toxification are yet to be identified. Previously, analysis of the genome of lab strain of *M. xanthus* reveled presence of 19 biosynthetic clusters that are predicted to be involved in the synthesis of diffusible antimicrobial substances including antibiotics^45^. Here we demonstrate that with respect to the abundance of antibiotic resistant bacteria, *M. xanthus* associated soil communities are different from the ones that were not associated with *M. xanthus.* suggesting that in nature *M. xanthus* has a long-term influence on local microbial community dynamics. Therefore, a detailed study to identify the nature of diffusible substances involved will be crucial for two reasons. First, understanding of the mechanisms driving community structure in the presence of *M. xanthus*. Second, to decipher the molecular basis for the observations reported in this manuscript.

Spread and abundance of antibiotic resistance has been primarily linked to the anthropogenic activities. However, it is becoming increasingly evident that resistance alleles can evolve even in the absence of antibiotics in the environment^46–48^. Though our results do not demonstrate the evolution of antibiotic resistance, they demonstrate the possibility that microbial interactions can influence resistance in natural communities. Recently it was demonstrated that social interactions can modulate mechanisms and rate of emergence of antibiotic resistance in the presence of antibiotics^49^. These results demonstrated that obligate dependencies could reduce rate of emergence of resistance in cooperative populations. On the contrary, our results strengthen the possibility that microbes such as *M. xanthus* and other antimicrobial producers might accelerate the evolution of resistance in complex natural communities. Further, metagenome analysis in the past has also suggested that eukaryotic predators might also influence the abundance of antibiotic resistance bacteria, but it is unclear whether the predators themselves are the causal agents or it is in fact the bacterial interactions that drive the changes in the frequency of resistant isolates^50^. Therefore, we call for a broader investigation which will increase our awareness of the influence of microbial interaction dynamics on the ecology and evolution of resistance mechanisms. These studies are crucial for the overall understanding of the spread of antibiotic resistance in nature, and especially in pristine environments.

## Acknowledgements

The authors thank Jessica M. Laloo from Bacterial Ecology and Evolution group, Department of Microbiology and Cell Biology at IISc, Bangalore, for her help with the illustrations in figures S4 and S5. The authors also thank DST-FIST for support to the Department of Microbiology and Cell Biology, UGC Centre for advanced study, and the DBT-IISc partnership. This study was supported by the funds from India Alliance Intermediate fellowship to SP (IA/I/20/1/504921).

## Author contribution

SP conceived the project. TSS made the initial observation, VM independently confirmed the initial observation. SS, JK and SP designed experiments. SS performed the experiments and JK analysed the data. SP, SS, and JK designed the metagenome analysis. JK analysed the metagenome data with the help of SW, SP and SZ. SP and SS wrote the first draft of the manuscript. SP edited the manuscript. All authors amended the manuscript.

## Methods

### Strains and culture condition

*M. xanthus* isolate S2, S3 and CVH1 were isolated from the Indian Institute of Science (IISc) campus using methods described in Kraemer & Velicer, 2011^51^. Whereas isolate MC2, MC8 and GH1 were previously reported as MC3.5.9_C5 and MC3.5.9_C29, and GH3.5.6_C27 respectively (reported previously in Pande et al^38^). For isolation of natural isolates, soil was collected with the help of a sterile 10 mL syringe from which the tip had been removed. After removing 5 mm of soil from each end of the column rest of the core was crushed and plated on a selective agar medium [CTT^52^ medium with 1.5 % bacto agar, Difco), containing antibiotics and antifungals, Vancomycin (10 mg/L, Sigma), Nystatin (1000 units/L, Sigma), Cyclohexamide (50 mg/L, Sigma ) and Crystal violet (10 mg/L, Sigma)]. Plates were incubated at 32 °C for two weeks. Following incubation plates were examined for presence of fruiting bodies on the soil surfaces. Single fruiting body was picked with a sterile toothpick and transferred to 1 mL autoclaved distilled water, incubated at 50 °C for 2 h, sonicated, and plated on CTT soft agar medium (0.5 % bacto agar, Difco). After 6 days of incubation at 32 °C clones were randomly picked. *M. xanthus* strains were stored frozen in liquid CTT^53^ medium with 20 % glycerol at -80 °C.

Three different media types were used to culture and estimate non-myxobacterial isolates directly in natural soil communities, and in complex soil communities cultured in the lab. Two types of LB medium i.e., ten times diluted (0.1 x) and standard LB media (HiMedia), and ten times diluted (0.1 x) TSB (Tryptic Soy Broth, HiMedia) medium. CTT medium was used to grow and estimate the population size of *M. xanthus*. Except for the experiments conducted in natural locations all experiments were conducted at 32 °C, and liquid cultures were incubated in shaking (200 rpm) conditions.

### Generation of S3 Rif strains

*M. xanthus* is naturally resistant to gentamycin and sensitive to salt. Therefore, CTT medium with Rifampicin has been used in many studies to distinguish *M. xanthus* from cocultured prey species. Similarly, media with salts (such as LB and TSA (Tryptic Soy Agar)) can be used to estimate population size of non myxobacterial isolates. However, since soil samples harbored Rifampicin resistant non-myxobacterial isolates that can grow on CTT, we generated Rifampicin resistant variant of S3 isolate. Dual resistance coupled with distinct colony morphology allowed us to estimate population size of *M. xanthus* in mixed culture experiments.

To generate Rifampicin resistant variant isolate S3 was grown in 8 mL CTT medium in a 50 mL flask for 24 h, at 32 °C, 200 rpm. After 24 hours, the culture was diluted into eight 50 mL flask with 8 mL CTT media, incubated at 32 °C at200 rpm, and was grown up to 0.5-0.6 OD. The complete content of each flask was plated in CTT soft agar with Rifampicin (5 µg/mL, MP Biochemicals) in eight 150 mm plates, respectively. The plates were incubated for two weeks, and the eight Rifampicin resistant colonies were picked. Resistance to antibiotics can impart some cost of resistance and hence influence the growth with respect to the parental strain. Hence, the growth neutrality of the Rifampicin resistant strains was tested by growing both the parental (S3 strain) and Rifampicin resistant (S3 Rif strains) in CTT liquid till mid-log phase, adjusting their density to 5x10^9^ cells/mL and spotting 10 µL of this density adjusted cultures on 20 mL CTT hard agar (1.5 % bacto agar) in 90 mm plates. The relative swarms of the strains on agar beds were recorded as a parameter of determining their respective growth. The relative growth of the resistant strain was to that of its parent stain was estimated by taking a ration of their swarm on CTT hard agar. For further experiments, S3Rif4 strain was selected since the relative growth of this strain was statistically indistinguishable from its parent strain (Data not shown here).

### Estimation of antibiotic resistance frequency

To estimate frequency of resistant isolates samples were dilution plated on antibiotic free 0.1x TSA (Tryptic Soy Broth, HiMedia + 0.5 % Agar-Agar, Qualigens) media and on 0.1x TSA media with one antibiotic for which the resistance frequency was to be estimated. We used six different antibiotics for the experiments (Kanamycin (40 µg/mL, Sigma Aldrich), Rifampicin (5 µg/mL, MP Biochemicals), Ampicillin (50 µg/mL, Sigma), Cycloheximide (5 µg/mL, Sigma), Gentamycin (10 µg/mL, SRL) and Vancomycin (1 µg/mL, Sigma) and Tetracycline (10 µg/mL, Sigma). Plates were incubated at 32 °C for 2 days.

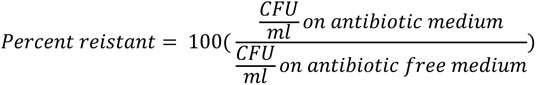

### Sample collection and classification of natural soil communities to test correlation between presence of *M. xanthus* and resistance in nature

Soil from twenty-five randomly selected locations was used to estimate the frequency of antibiotic resistance in nature and its correlation with the presence or absence of *M. xanthus*. In brief, soil samples were collected with the help of a sterile 10 mL syringe from which the tip had been removed. After removing 5 mm of soil from each end, central core from the soil column was equally split to test for the presence of *M. xanthus* and to estimate the frequency of resistant isolates.

To estimate the frequency of antibiotic resistant isolates 4 mL of each soil column was dissolved in 20 mL of sterile water and mixed thoroughly. Next, the soil samples were allowed to stand for 4 h at room temperature, and the antibiotic resistance frequency assay (See estimation of antibiotic resistance frequency) was performed with the natural soil communities obtained from the supernatant of the soil samples. In our experience 4 h incubation allowed for highest CFU count in the conditions used in our experiments, suggesting most microbes dissociated from soil particle. No technical replicates for the estimation of the frequency of resistant bacteria in the soil sample were performed, and therefore each estimation was used as a single biological replicate for statical analysis.

Each soil sample was also tested for the presence of *M. xanthus*. For this 4 mL soil core from each sample was crushed and plated on selective media [modified CTT medium with 0.5 % casitone, 1.5 % agar, Vancomycin (10 mg/L), Nystatin (1000 units/L), Cyclohexamide (50 mg/L) and crystal violet (10 mg/L)]. *M. xanthus* is a gram-negative soil bacterium that forms spore filled multicellular fruiting bodies upon starvation. These fruiting bodies are easily visible under light microscope, and hence provide a simple mode of detection for Myxobacteria in natural soil isolates. We used presence or absence of fruiting bodies on selective media to classify each soil sample either as *M. xanthus* positive (M+) or negative (M-), by visually observing for presence (designated M+) or absence (designated M-) of *M. xanthus* fruiting bodies.

### Influence of *M. xanthus* on antibiotic resistome in natural soil communities in laboratory conditions

Microbial communities were derived from four random locations (named Location-1, Location-2, Location-3, and Location-4) inside IISc. campus. The soil samples were collected up to the mark of 10 mL in sterile 50 mL falcon tubes. To extract the communities from these derived soil samples, 20 mL of sterile double distilled water was added to these soil samples, mixed uniformly, and allowed to stand at room temperature for 4 h. Once most of the sediments had settled down, 100 µL of these communities (supernatant post-sedimentation) were inoculated with or without 1 mL (10^6^ cells) of Rifampicin resistant S3 strain, in 8mL of 0.1x TSB liquid in 50 mL flasks. The cultures were incubated at 32 °C, 200 rpm for 6 days.

Resistance frequency was measured by dilution plating cultures 0.1x TSA + 0.5 % Bacto agar (soft agar medium) either supplemented with or without different antibiotics (See *estimation of antibiotic resistance frequency)* over a period of 6 days, with plating done every 24 h, starting from 0 h. To estimate the growth of *M. xanthus* within these communities, dilution plating was done in CTT 0.5% agar medium with Rifampicin (final concentration 5 µg/mL) and Gentamycin (final concentration 10 µg/mL every 24 h over a period of 6 days. These experiments were performed in three independent blocks of biological replicate for each of the soil sample.

### Influence of spent media from *M. xanthus* supplemented communities on antibiotic resistome in natural soil communities

Spent media was collected from three distinct soil communities cultured in laboratory condition, either supplemented with or without *M. xanthus*, as described earlier (See Influence of *M. xanthus* on antibiotic resistome in natural soil communities in laboratory conditions). The spent media was collected by sampling the cultures every 24 h and spinning down the cultures to extract the supernatant. The supernatant collected from these samples were filter sterilised using 0.20 -micron membrane filter (Minisart, Sartorius Stedim Biotech GmbH, Germany) to obtain cell-free spent media. To further estimate whether the presence of diffusible molecules in the spent media were responsible for the enrichment of resistant isolates, freshly collected soil communities were subjected to the spent media extracted from communities where either *M. xanthus* were either added or not added (as controls), diluted with fresh 0.1x TSB liquid media (1:1). 100 µL of freshly isolated communities were further cultured in the spent media from the respective two conditions, for 6 days at 32 °C and 200 rpm. Post-incubation the frequency of resistant isolates was estimated by dilution plating as described earlier (See estimation of antibiotic resistance frequency). These experiments were performed in three independent biological replicates.

### Influence of spent media from *M. xanthus* supplemented community on growth of common prey bacteria found in soil

Though the introduction of spent media enriched the overall resistant isolates in natural communities, to further understand whether this enrichment is the result of growth inhibitory effects of the diffusible molecules present in the spent media, that were extracted from cultures inoculated with *M. xanthus*, common prey species were grown under laboratory culture conditions either in presence or absence of the spent media. Five distinct prey species were used for this assay, *Arthrobacter globiformis, Escherichia coli, Pseudomonas putida* and *Rhizobium vitis.* These preys were grown to mid-log phase in LB liquid and OD was adjusted to 0.1 OD using buffer. 100 µL of the density adjusted cultures was inoculated in 8 ml of media supplemented with either the cell-free spent media or buffer in 1:1 ratio. The cultures were incubated at 32 °C, 200 rpm for 24 h. The final ODs were recorded at 600nm to determine the growth of different bacteria (thermoScientific, Varioskan LUX). These experiments were performed in three independent biological replicates.

### Influence of *M. xanthus* lysate on resistome of natural communities

To understand if the enrichment of resistant isolates is brought about specifically as a result of *M. xanthus’* lysis, lysate of *M. xanthus* (S3Rif strains) cells was prepared by removing the growth media, resuspending the exponentially growing vegetative cells in buffer and sonicating (25 % intensity, 10 cycles of 30 sec ON and 10 sec OFF, Qsonica-700 sonicator) them to lyse the vegetative cells. 4 mL of this lysate was further introduced along with 100 µL of freshly isolated soil communities in 4 mL of 0.1x TSB liquid, under previously described culture conditions for 6 days. As a control, communities were also cultured under similar condition without the addition of the lysate. The final resistance frequency was recorded by dilution plating on 0.1x TSA with and without different antibiotics (See estimation of antibiotic resistance frequency). These experiments were performed in three independent biological replicates.

### Influence of supernatant extracted from *M. xanthus* fruiting bodies on resistome of natural communities

*M. xanthus* (S3Rif strains) cells were grown in CTT liquid to mid-log phase and the cell-density was adjusted to 5x10^9^ cell/mL. 100 µl of this density adjusted cultures were spotted on 10 mL of starvation media in 60 mm plates (TPM media, same composition as that of CTT without the casitone as carbon source + 1.5 % Bacto agar). The plates were incubated for 3 days at 32°C to allow the development of fruiting bodies. After 3 days incubation, the fruiting bodies was harvested by washing the TPM agar beds with 1 mL of TPM buffer and the supernatant was extracted after spinning down the resuspended spores. The supernatant was further filter sterilised using 0.45-micron membrane filter. 100 µL of freshly isolated soil communities were then cultured in 8mL of 0.1x TSB liquid with 1 mL of supernatant extracted from the fruiting bodies. As controls, 1mL of TPM buffer was added to the communities. The culture conditions were similar to previous experiments, and the cultures were incubated at 32 °C, 200 rpm for 6 days. Post 6 days incubation, the resistance frequencies were determined by dilution plating on 0.1x TSA with and without different antibiotics (See estimation of antibiotic resistance frequency). These experiments were performed in six independent biological replicates.

### Sample preparation and DNA extraction for metagenome sequencing

Soil sample of three independent locations were collected and communities were derived as described above (See Influence of *M. xanthus* on antibiotic resistome in natural soil communities in laboratory condition*s*). The communities were cultured in 8 mL of 0.1x TSB with or without *M. xanthus* as described earlier. Cultures incubated for 6 days were sent to sequencing facility for isolation and shotgun metagenome sequencing. DNA extraction and shotgun metagenome sequencing was done by the sequencing agency, Nucleome Informatics, Hyderabad, India. For isolating the total DNA from the communities, CTAB method was used. To 50 µL of the culture sample 700 µL of CTAB buffer was added, vortexed briefly and incubated at 65 °C for 45 mins. Post incubation, the DNA was purified using phenol-chloroform extraction method and eluted in nuclease free water. Primary QC was performed with the extracted DNA using Nanodrop and gel. Finally, genomic libraries were prepared using KAPA HyperPlus Kit (cat no-KR1145-v8.21). Sequencing was done using Illumina Novoseq 6000 S4 flow cell.

### Influence of *M. xanthus* on antibiotic resistome in nature

Natural soil samples were collected from 15 locations within the *IISc* campus. From each location, soil was collected up to the mark of 10 mL in sterile 50 mL falcon tubes. An additional amount of soil was collected from each of these locations, which was autoclaved, and re-used as sterile soil samples for respective locations. Next, 20 mL water was added to each falcon, thoroughly mixed, and kept standing in room temperature for 4 hours. 1 mL soil supernatant with 1 mL (10^6^ cells) Rifampicin resistant variant of S3 isolate was added to previously autoclaved 20 mL soil samples derived from respective location. Falcons were mixed by inverting for 12 times, planted at their respective locations, and incubated for 5 days. For these experiments we constructed 50 mL falcons which only allow exchange of resources (but not cell) between internal and external environment (Figure S4). In control experiments soil supernatant without S3 isolate was inoculated in respective soil sample.

To harvest the communities within the soil sample, after 5 days of incubation, 20 mL of autoclaved water was added to each falcon, mixed thoroughly and allowed to stand for 4 h at room temperature. 100 µL supernatant from each falcon was dilution plated on 1x LB and 0.1x LB (diluted LB media) agar plate (0.5 % agar) with or without antibiotics, incubated at 32 °C. For this, we used six different antibiotics namely, Kanamycin (40 µg/mL), Rifampicin (5 µg/mL), Ampicillin (50 µg/mL), Cycloheximide (5 µg/mL), Gentamycin (10 µg/mL) and Vancomycin (1 µg/mL). *M. xanthus* cells do not grow in LB medium. Therefore, growth of *M. xanthus* was measured by dilution platting 100 µL supernatant in CTT 0.5 % agar with Rifampicin and Gentamycin (*M. xanthus* is naturally resistant to Gentamycin). These experiments were performed in three independent biological replicates.

### Metagenome data processing

Metagenomic reads were processed using the metagenome-atlas workflow v2.18.1^53^ with default parametrisation if not stated otherwise in the following description. In brief, the applied atlas workflow consisted of three main steps: (1) Quality control and filtering, (2) read assembly, and (3) binning of contigs. For step (1 – quality filtering), reads were quality trimmed and reads, which are likely contaminations from Illumina PhiX control sequences, were removed using functions from the BBmap suite v39.01 (BBMap - Bushnell B. - sourceforge.net/projects/bbmap/). In step (2), reads were assembled into contigs using metaSPAdes v3.15.5^54^. For the binning step (3), assembled contigs were assigned to bins using metabat v2.15^55^. The completeness and contamination percentages of refined bins were estimated using checkM2 v1.0.1^56^.

Subsequently, filtered bins were clustered, and cluster-representative bins were selected using dRep v3.4.5^57^ by re-using the contamination and specific completeness percentages predicted by checkM2 from the previous analysis step. Secondary clustering was performed for the average nucleotide identity of >= 95% as threshold. Representative bins (from here on termed MAGs) were quantified in each metagenome sample by mapping quality control-filtered reads to the MAG genomic sequences using coverM v0.6.1^58^ in default parametrisation. Taxonomic classification of MAGs was predicted using GTDB-tk v2.3.2^59^ and the reference data version r207_v2.

### ARG analysis

Technical University of Denmark (DTU) sponsored KmerResistance ARG tool (v2.2.0)^60,61^ was used to estimate antibiotic resistance alleles. KmerResistance is available at http://genomicepidemiology.org/services/. Clean reads’ fastq files for day 6 were submitted to KmerResistance algorithm which were then mapped based on the co-occurrence of k-mers against a class of resistance alleles that were available in ResFinder database. A minimum of 90% coverage, and 90% query identity cut-offs were used to determine the best matchable resistance alleles. The identified alleles were then grouped into their respective resistance classes such as aminoglycoside, betalactam, fosfomycin.

### Statistical Analysis

All data analysis was done using R (Version 4.3.1). Each experiment was performed in at least three or more blocks of independent replicates unless otherwise specified. Data was checked for homogeneity and appropriate statistical tests were used. Wherever required, multiple testing was corrected for false discovery rate. Wherever applicable, data transformations are mentioned in figure legends.

