## Supplementary information for "Mass lysis of bacterial predators drives the enrichment of antibiotic resistance in soil microbial communities"

**Supplementary figure 1:**

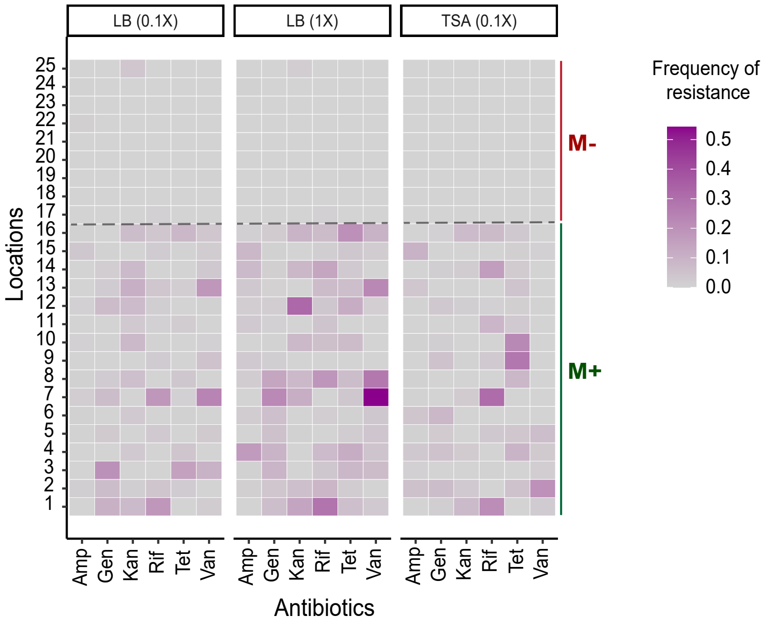


**Figure S1: Distribution of antibiotic resistance across soil types (M+/M-) in different media conditions.** Out of the 25 soil samples collected, 16 were classified as *M. xanthus* positive (M+) based on visual detection of *M. xanthus* fruiting bodies (See methods) and 9 were classified as M. *xanthus* negative (-M). The overall antibiotic resistance frequency was simultaneously checked in all 25 soil samples across 6 distinct clinically relevant antibiotics (Ampicillin, Gentamicin, Kanamycin, Rifampicin, Tetracycline and Vancomycin). Estimation of resistance frequency was performed in three different media conditions, 0.1xLB, 1xLB and 0.1x TSA to allow the growth of a broad range of culturable bacteria. The heat-map shows the relative frequencies of antibiotic resistance across different antibiotics are generally higher in M+ soil samples. Darker colour implies a higher frequency of resistant bacteria.

**Supplementary figure 2:**

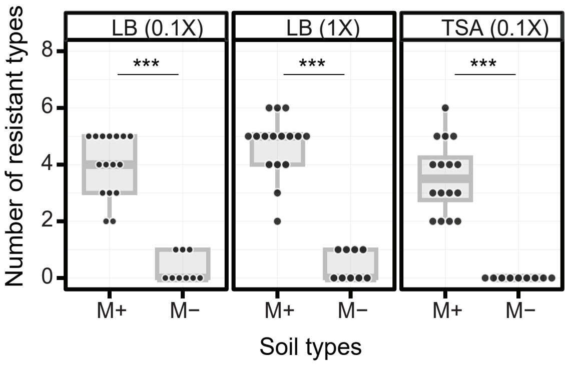


**Figure S2: Number of antibiotics against which resistance was observed in three culture conditions, 0.1x LB, 1x LB, 0.1x TSA.** Each dot represent the number of antibiotics against which resistance was observed within a single soil sample (Independent-sample t-test between M+ and M- conditions in each media type, LB (0.1X): p = $3.864e^{-12}$; LB (1X) : p = $2.2e^{-16}$; TSA (0.1X) : p = $1.002e^{-08}$, n = 3).

**Supplementary figure 3:**

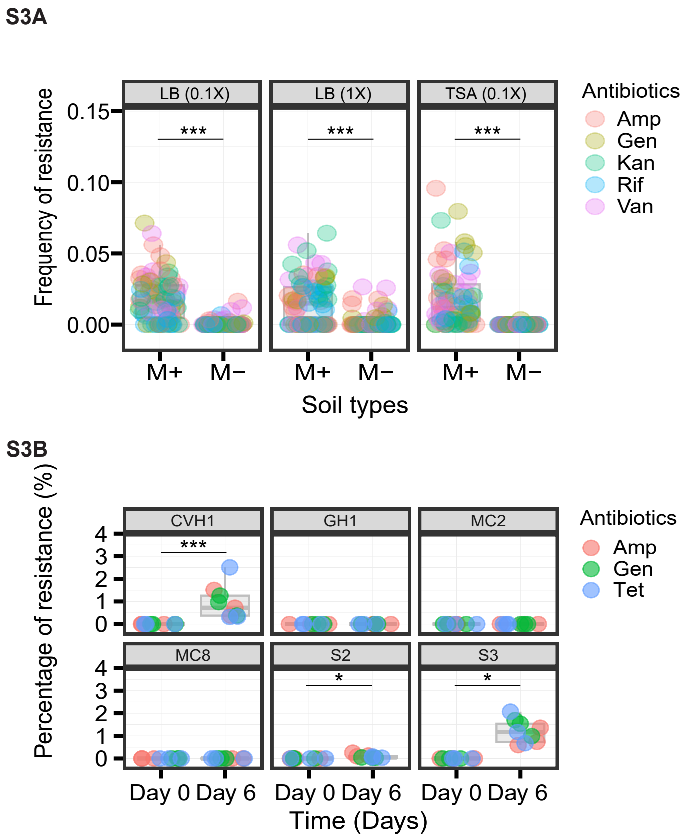


**Figure S3: Effect of culture conditions and distinct natural *M. xanthus* isolates on the enrichment of antibiotic resistance.** (A) Frequency of resistant isolates is enriched on addition of *M. xanthus* (M+) compared to when *M. xanthus* is not added (M-). Data shown is for resistance frequencies detected in M+/M- conditions across three distinct media types. Each dot represents antibiotic frequency for one antibiotic in one soil community (total 4 communities were tested). (Paired-sample t-test between M+ and M- conditions in each media type, LB (0.1X): p = $4.218e^{-07}$; LB (1X) : p = $5.554e^{-07}$; TSA (0.1X) : p = $4.467e^{-}05$, n = 3). (B) Frequency of resistant isolates on $0^{th}$ day and after 6 days of incubation with different *M. xanthus* isolates (CVH1, GH1, MC2, MC8, S2, S3) is shown. Each dot represents antibiotic frequency for one antibiotic in one soil community (4 communities were tested) (Paired-sample t-test between Day 0 and Day 6 for addition of each *M. xanthus* isolate, p < *0.05*, n = 3).

**Supplementary figure 4:**

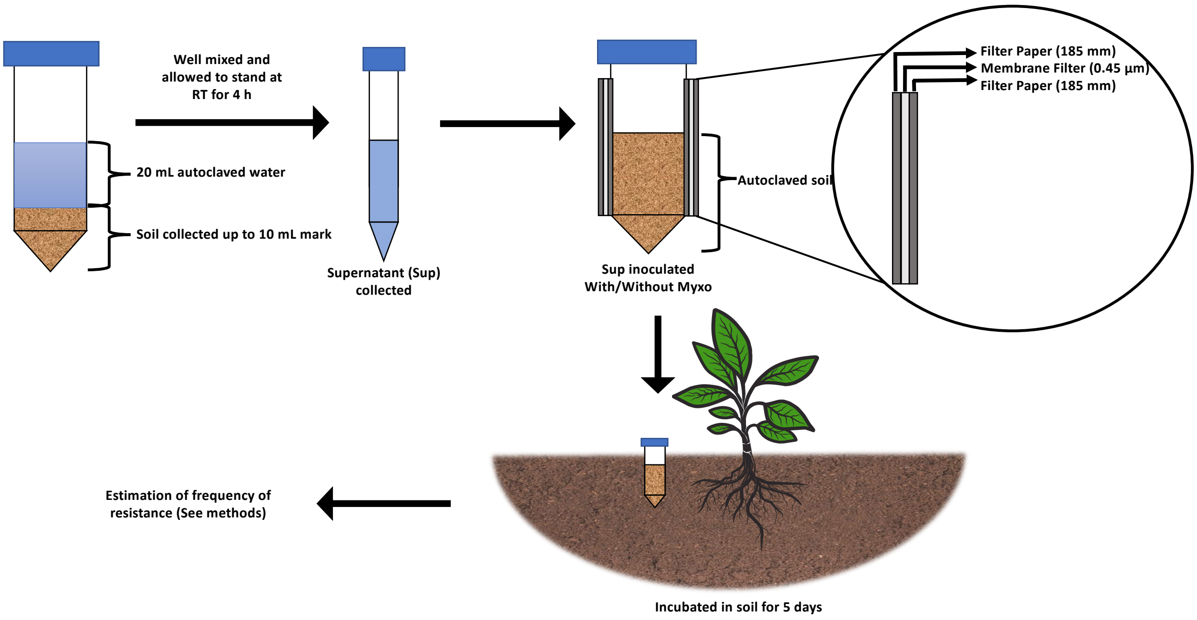


**Figure S4: Schematic representation of experimental design.** Soil communities were isolated by adding autoclaved water to the collected soil sample in a 50 mL tube, soil was mixed with water by shaking, and the samples were allowed to stand at room temperature for 4 h. This allows most of the soil sediments to settle down and the communities are extracted by collecting the supernatant. The isolated soil supernatant were then further inoculated with or without *M. xanthus* in autoclaved soil in a modified 50 mL tube, whose sides were cut off and sealed with triple layer of filter paper-membrane (0.4 micrometer pore size). This modification allowed diffusion of molecules from the external environment but prevented any further contamination of soil sample. These tubes were then buried in the respective locations, from where the soil samples were isolated. After 5 days of incubation, the communities were extracted from soil samples in the laboratory and antibiotic resistance frequencies were estimated (See methods)

**Supplementary figure 5:**

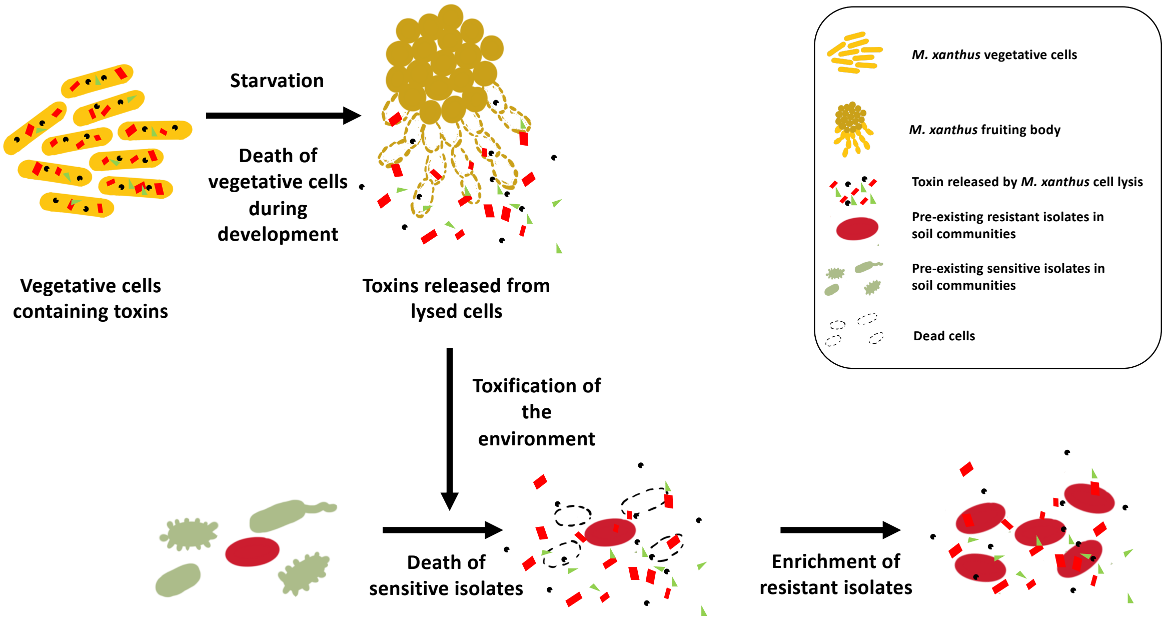


**Figure S5: Diffusible substances released during fruiting body development by *M. xanthus* cells can enrich resistant isolates in soil communities.** Starvation induced fruiting body formation and development is an essential life-history trait of *M. xanthus*. During development, almost 90 % of the *M. xanthus* population die. We demonstrate the resistance enriching effect of the supernatant extracted from starved *M. xanthus* population that was allowed to fruit for 3 days (Figure 5B). This is in addition to the supernatant enriching effect of M. xanthus cell lysate (Figure 4C). Therefore, we hypothesize, that in soil, where nutrient conditions are expected to fluctuate, cell lysis during fruiting body formation can toxify the environment, which can then result in the enrichment of overall antibiotic resistance. Together these observations explain the correlation between presence of *M. xanthus* with higher frequency of antibiotic resistant bacteria.
